# Female fruit flies cannot protect stored sperm from high temperature damage

**DOI:** 10.1101/2021.09.30.462530

**Authors:** Benjamin S. Walsh, Steven R. Parratt, Rhonda R. Snook, Amanda Bretman, David Atkinson, Tom A. R. Price

## Abstract

Recently, it has been demonstrated that heat-induced male sterility is likely to shape population persistence as climate change progresses. However, an under-explored possibility is that females may be able to successfully store and preserve sperm at temperatures that sterilise males, which could ameliorate the impact of male infertility on populations. Here, we test whether females from two fruit fly species can protect stored sperm from a high temperature stress. We find that sperm carried by female *Drosophila virilis* are almost completely sterilised by high temperatures, whereas sperm carried by female *Zaprionus indianus* show only slightly reduced fertility. Heat-shocked *D. virilis* females can recover fertility when allowed to remate, suggesting that the delivered heat-shock is destroying stored sperm and not directly damaging females in this species. The temperatures required to reduce fertility of mated females are substantially lower than the temperatures required to destroy mature sperm in males, suggesting that females are worse than males at protecting mature sperm. This suggests that female sperm storage is unlikely to ameliorate the impacts of high temperature fertility losses in males, and instead exacerbates fertility costs of high temperatures, representing an important determinant of population persistence during climate change.

## Background

Anthropogenic climate change poses a significant challenge to global biodiversity. We urgently need to understand how rising average temperatures, and an increasing number of short-term extreme temperature events (Perkins-Kirkpatrick and Lewis, 2020), will affect natural populations. Understanding how high temperatures affect organisms can allow researchers to predict the vulnerability of species and inform conservation efforts, revealing which temperature-sensitive traits are particularly important for determining species persistence. Initial research focused on temperatures required to kill individuals, and it has been shown that species’ lethal temperatures correlate with the maximum temperatures species experience in the wild (Kellermann et al., 2012). It has been known for around a century that high temperatures can sterilise individuals (Cowles, 1945; David et al., 2005; Young and Plough, 1926). Recent work has found that the temperature that sterilises over 80% of males in a species, named a species’ upper thermal fertility limit (TFL), correlate more strongly with maximum temperatures that species experience in the wild (Parratt et al., 2021; van Heerwaarden and Sgrò, 2021). This indicates that upper TFLs are significant determinants of current species distributions, and are therefore likely to shape population persistence as climate change progresses.

Temperature-induced sterility occurs across a wide-variety of taxonomic groups (David et al., 2005; Hurley et al., 2018; Karaca et al., 2002; Sage et al., 2015; Walsh et al., 2019b). A study of 43 *Drosophila* fruit fly species found that males from nearly half the species (19/43) are sterilised at temperatures significantly lower than temperatures required to kill them (Parratt et al., 2021). Male fertility generally seems more sensitive to high temperatures when directly compared with female fertility (Iossa, 2019; Sales et al., 2018; Walsh et al., 2020), although the converse is possible (Janowitz and Fischer, 2011). The relative sensitivity of male fertility in animals has been attributed to disruption of spermatogenesis or death of mature sperm as a result of thermal stress (Rohmer et al., 2004; Sales et al., 2018). Typically, the effect of temperature on fertility is measured by directly heating males, and subsequently measuring the reproductive capacity of focal males when paired with females following heat-stress (Jørgensen et al., 2006; Karaca et al., 2002; Parratt et al., 2021; Sales et al., 2018; Walsh et al., 2020; Zwoinska et al., 2020) or by measuring other traits linked to fertility (Hurley et al., 2018; Paxton et al., 2016). Likewise, studies measuring female fertility generally stress females prior to mating (Sales et al., 2018; Walsh et al., 2020; Walsh et al., 2019a), in order to isolate the effect of temperature on female reproductive physiology, such as oocytes. However, while it is clearly important to measure the effect of thermal stress prior to mating, the effect of high temperatures on females post-mating has been largely ignored. This is important because sperm can spend a significant proportion of time within the female reproductive tract prior to fertilisation.

Sperm storage is characterised by temporal delays between insemination and fertilisation, during which sperm is maintained within a female’s reproductive tract. Female sperm storage is common across taxa, including mammals, birds, reptiles, fish and insects (Holt, 2011; Sever and Hamlett, 2002). The time that sperm can be kept viable inside a female varies substantially. In birds and reptiles, sperm storage durations range from seven days up to seven years, in mammals for less than a day up to six months in some bat species, amphibians from four to thirty months, in fish from only days to around two years, and over a decade in some eusocial hymenoptera (Birkhead and Møller, 1993; Holt and Lloyd, 2010; Holt, 2011; Keller, 1998; Levine et al., 2021; Pamilo, 1991). The method of sperm storage can also vary substantially, and phylogenetic evidence suggests long-term storage of sperm has arisen independently across taxa (Holt and Lloyd, 2010). For example in birds and some reptiles, inseminated spermatozoa are stored in microscopic sperm storage tubules (SSTs) embedded in the infundibulum, which allow sperm to survive for extended periods of time (Holt, 2011; Sasanami et al., 2013). Females from the majority of insects and some other arthropods store sperm in a highly chitinised specialised organ called the spermatheca. Most insects have one spermatheca, but some insects have two or three (Pascini and Martins, 2017). However, while female sperm storage for extended durations is taxonomically widespread (Birkhead and Møller, 1993), the impact of high temperatures on sperm stored within mated females is currently understudied. The few efforts to examine the impact of high temperatures on sperm stored within females include mated females of the red flour beetle (*Tribolium castaneum*), which show a 33% reduction in offspring production when exposed to a heatwave treatment (Sales et al., 2018). Also, a four hour heat-stress at 42°C significantly reduces the viability of sperm stored by honey bee queens (McAfee et al., 2020), although in this study the authors do not directly test whether this reduces female offspring production. Given the urgency of understanding the consequences of rising temperatures, we need a better understanding of the thermal robustness of female sperm storage.

Fruit flies from the family Drosophilidae provide a useful model group to explore this question. Female *Drosophila* typically possess a pair of spermathecae and a seminal receptacle, the latter of which is a thin extended tubule arising from the uterus (Pitnick et al., 1999). *Drosophila* have been proposed as a model system for studying sperm-female interactions, in order to better understand fertilisation across taxa (Heifetz and Rivlin, 2010). *Drosophila* are also a model taxon for studying thermal reproductive physiology, including examining how high temperatures affect fertility of both males and females prior to mating (David et al., 2005; Parratt et al., 2021; Sgrò et al., 2016; Walsh et al., 2020). However, to our knowledge there has been no substantial effort to examine how high temperatures affect the capacity of mated females to produce offspring in *Drosophila*.

Here, we explore the impact of heat stress on sperm storage in females from two *Drosophila* species. We test the tropical pest species *Zaprionus indianus*, and a more temperate species *Drosophila virilis*. Parratt et al. (2021) showed that males of both species die when exposed to ∼38°C for 4 hours, and(Parratt et al., 2021) that mature sperm are destroyed at ∼37°C for 4 hours when stored in male *D. virilis*, but not in male *Z. indianus*. In contrast, the same study found that developing sperm appear to be destroyed by high temperatures in both species. Males of both species are sterile 7 days after being heated at ∼35°C for 4 hours. However, we do not know the effect of high temperatures on sperm stored within mated females.

We test three components of female fertility across time. Firstly, we test the expectation that female fertility will be more robust to high temperatures than male fertility. Secondly, we test whether sperm stored in mated females are more or less sensitive to high temperatures than sperm stored in a male’s seminal vesicles and developing sperm within the testes, investigated previously. Finally, we explore whether mated females that are heated to a point that sterilises them can recover fertility, after being presented with new male partners. If sterilised mated females can recover by remating, this would suggest that heat induced sterility of mated females is caused by damage to sperm and not direct damage to females.

## Materials and Methods

### Animal stock maintenance

Stocks of *Drosophila virilis* (Cambridge Fly Facility StrainvS-4, isolated in 1991) and *Zaprionus indianus* (DSSC Stock #: 50001-0001.05 ISOFEMALE, isolated in 2004), were kept in a temperature-controlled incubator (LMS 600NP Series 4) at 23°C, 12:12 L:D and ambient humidity. Stocks were maintained at moderate density (50 – 100 flies per 300ml bottle culture). *D. virilis* were kept on standard cornmeal-molasses-agar media, and *Z. indianus* were kept on banana medium. Ovipositing adults for both species were tipped to new food every week to prevent overlapping generations, and were replaced with fresh sexually mature adult flies every 4-6 weeks.

### Experimental treatments

Experimental treatments are summarised in Figure 1. We assessed whether heat stress influences fertility of females when delivered before mating (Experiment 1). We then completed an experiment with two more treatment combinations (Experiment 2a & 2b) to address the outstanding question of whether mated females can protect stored sperm from temperature damage experienced post-mating, and isolate effects on stored sperm from changes to female egg-laying behaviour.

**Figure 1:**
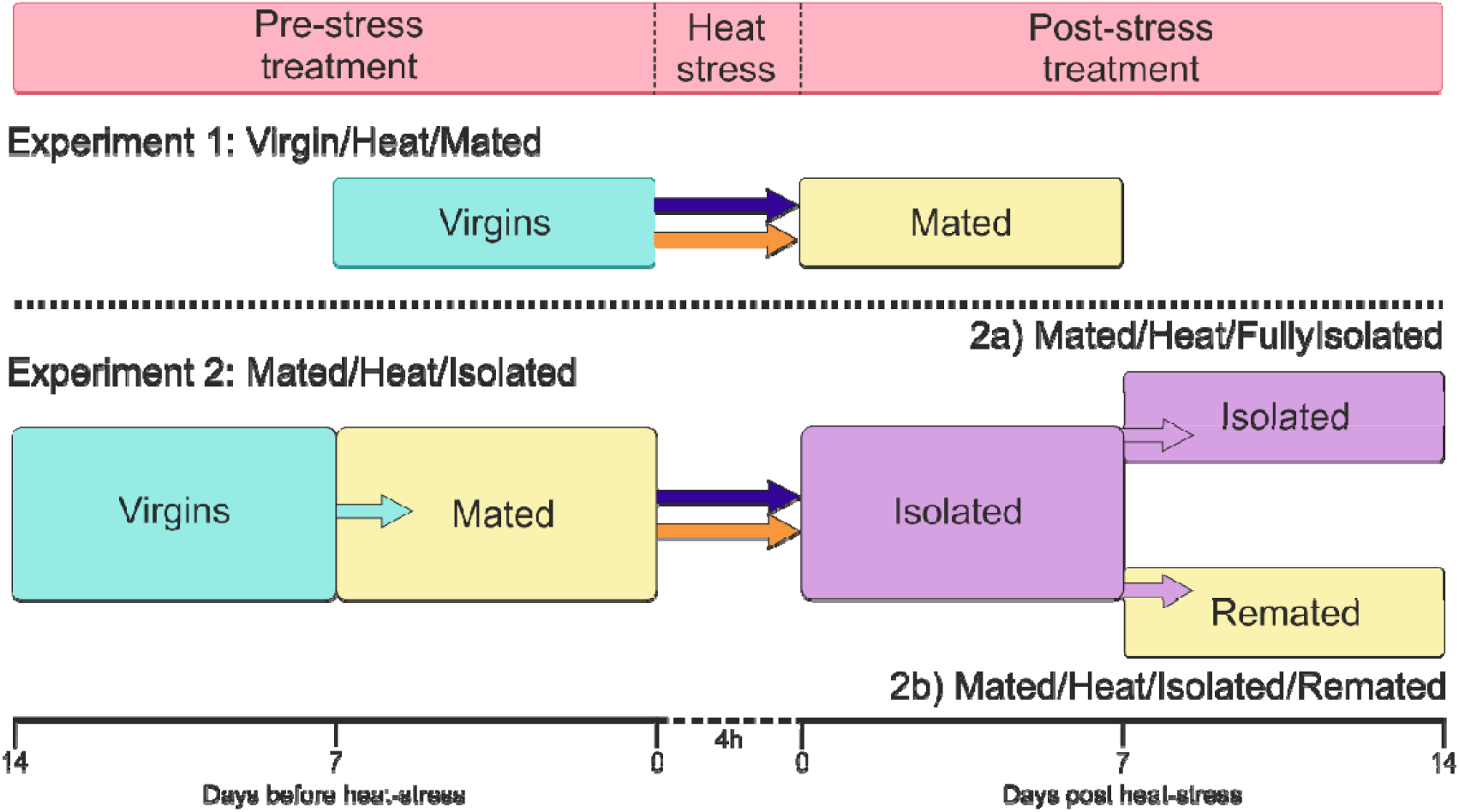
Experimental design outlining the two experiments. Each treatment designation combines various pre and post-stress mating treatments. **Experiment 1: Virgin/Heat/Mated**, where virgin females were heat-stressed and mated following heat-stress. **Experiment 2: Mated/Heat/Isolated**, where mated females are heat-stressed and kept alone for 7 days to produce offspring from previous matings. After 7 days post heat-stress, the experiment was divided into two treatments. For an additional 7 days, females were either kept in isolation **(2a: Mated/Heat/FullyIsolated)**, or given new male partners to mate with **(2b: Mated/Heat/Isolated/Remated)**. Focal females were exposed to either benign (23°C) or stress (35 & 36°C) temperatures for 4h in water baths. Day 0 in the post-stress treatment represents the time-point when the fertility assay begins (Fig. 2 & Fig. 3).

We chose to mate females at 7 days old when fully sexually mature, and kept this consistent between experiments. Therefore, females from Experiment 1 are 7 days old at heat-stress, whereas females from Experiment 2 are 14 days old at heat-stress. Prior to heat stress, females from Experiment 1 were separated at emergence and kept as virgins in groups of 10 for 7 days. Females from Experiment 2 were separated as virgins and kept in groups of 10 for 7 days, then provided with sexually mature males (7 days old) at a 1:1 sex ratio for a further 7 days prior to heat-stress. This produced an ‘assumed’ mated treatment, where females would have many opportunities to mate with a variety of males.

Immediately following heat stress, females were transferred to individual fresh food vials. In Experiment 1, virgin females were immediately placed with 4 virgin males. This mating group was moved to fresh vials twice, creating 3 ‘time-points’ where fertility was recorded. Females in Experiment 2 were isolated and transferred to fresh vials giving 3 time-points over 7 days. Experiment 2 was then split into two treatments. Females from Experiment 2a were kept in isolation for an additional 7 days, producing 3 more time-points where females were isolated. Females from Experiment 2b were placed with 4 males following the first 7 days of isolation. This mating group was transferred onto new vials twice more, giving 3 time-points where the females were isolated, followed by 3 recorded time-points where females were paired with males. Females were deemed as qualitatively fertile at a given time-point if there was evidence of larvae present in their vial (1/0), measured by directly observing larvae or their distinctive tracks in the food.

### Heat-stress

Groups of 10 females were transferred to fresh 25 × 95mm plastic vials, containing 25ml of ‘ASG’ medium (10g agar, 85g sucrose, 20g yeast extract, 60g maize, 1000ml H2O, 25ml, 10% Nipagin) to prevent desiccation. These vials were randomly assigned to pre-heated water-baths (Grant TXF200) for four hours at either control: 23°C, or two stress temperatures: 35°C & 36°C. The chosen temperatures do not affect survival or immediately sterilise mature adult males of either species, but result in substantial delayed sterility of males, likely due to the destruction of developing sperm (Parratt et al., 2021). Immediately following heat-treatment, flies were returned to benign temperatures (23°C).

### Statistical analyses

Species and experiments were analysed separately due to inherent differences in methodological design as summarised in Fig. 1. Treatment of females in Experiment 2 are identical from the start of the experiment until the experiment is split after 7 days into the post stress treatment. Therefore, data from Experiment 2 over the first 3 time-points were analysed together. The final 3 time-points of Experiment 2a after splitting were not statistically analysed, as all flies of both species in these final 3 time-points were completely sterile with only one exception, making these data uninformative. Experiment 2b was analysed after the treatments were split and females were presented with new males, in order to assess differences in fertility recovery across temperature treatments.

To assess the effect of temperature on fertility we used generalised linear mixed models with Bernoulli error distributions. We fitted fertility as a binary response variable, temperature and time-point and their interaction as fixed effects, and focal fly ID as a random effect to account for repeated measures. We did model selection using Wald Chi-squared likelihood ratio-tests, removing non-significant interactions. We retained all main effects and reported statistics of these from type II likelihood ratio tests using the ‘Anova’ function from the ‘car’ package, in the statistical software ‘R’. We then reported any pairwise comparisons in which p<0.05.

## Results

### Experiment 1: Virgin/Heat/Mated

There was no significant interaction between temperature and time on fertility of *D. virilise* from Experiment 1 (χ^2^_(2)_ = 3.977, p=0.137; Figure 2). There was no main effect of temperature (χ^2^_(2)_ = 0.093, p=0.954; Figure 2a), or time (χ^2^_(1)_ = 0.301, p=0.583; Figure 2) on fertility of *D. virilis*. Fertility was initially high, and remained so for the three time-points measured.

**Figure 2:**
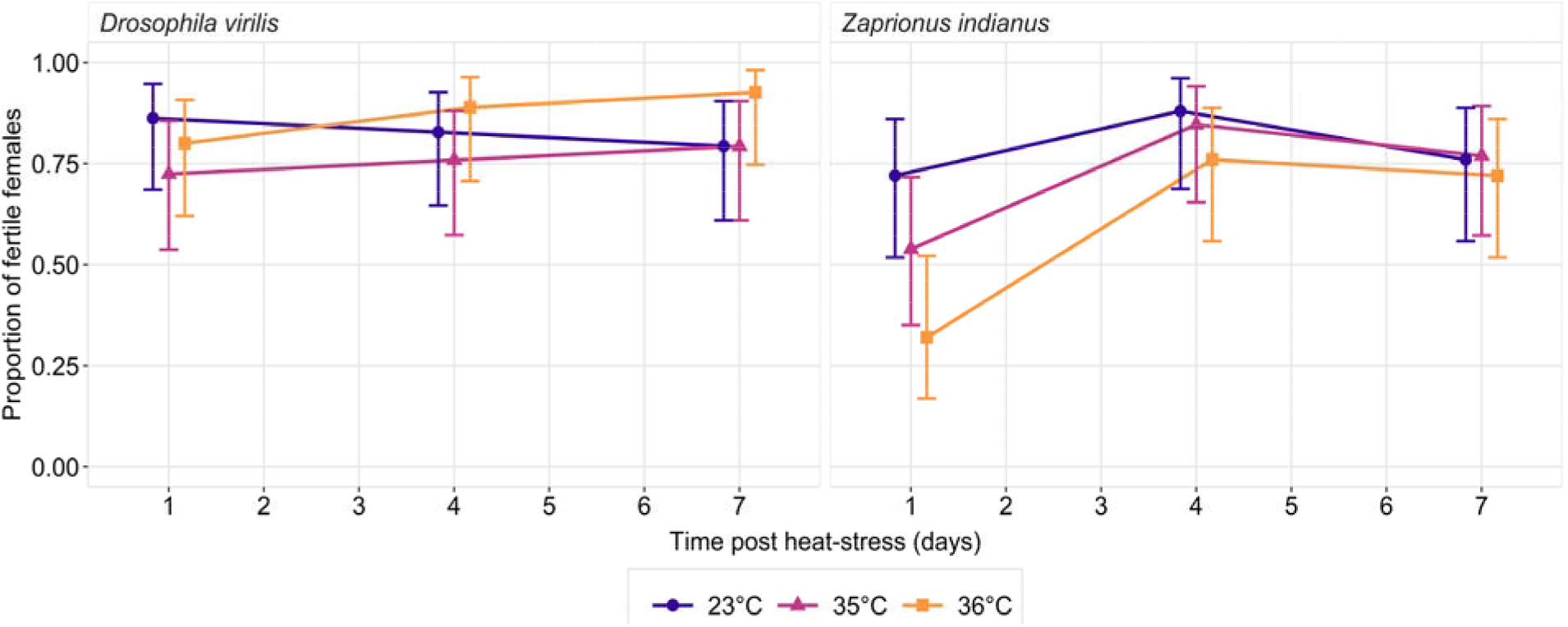
Proportion of fertile *D. virilis* and *Z. indianus* females over time for **Experiment 1: Virgin/Heat/Mated**, as described in Fig 1. Virgin females were heat-shocked at either benign (23°) or two stress temperatures (35 & 36°C) for 4 hours, and paired with 4 male partners immediately following heat-stress. This mating group was given 3 days to lay eggs, then tipped onto fresh vials twice, giving three recorded time-points where fertility was measured. Error bars represent 95% confidence intervals.

There was also no significant interaction between temperature and time on fertility of *Z. indianus* from Experiment 1 (χ^2^_(2)_ = 3.946, p=0.139; Figure 2). While the absolute proportion of fertile females heated at 36°C was consistently lower than controls, there was no overall main effect of temperature on fertility of *Z. indianus* (χ^2^_(2)_ = 4.469, p=0.107; Figure 2). However, there was a significant effect of time on fertility (χ^2^_(1)_ =10.911, p<0.001; Figure 2), where flies from all temperatures show increased fertility rates over time.

### Experiment 2: Mated/Heat/Isolated

Not all females from the pre-stress ‘mating’ treatment produced offspring, with controls producing a baseline fertility of around 70% in *D. virilis* and around 80% in *Z. indianus* (Fig. 3).

**Figure 3:**
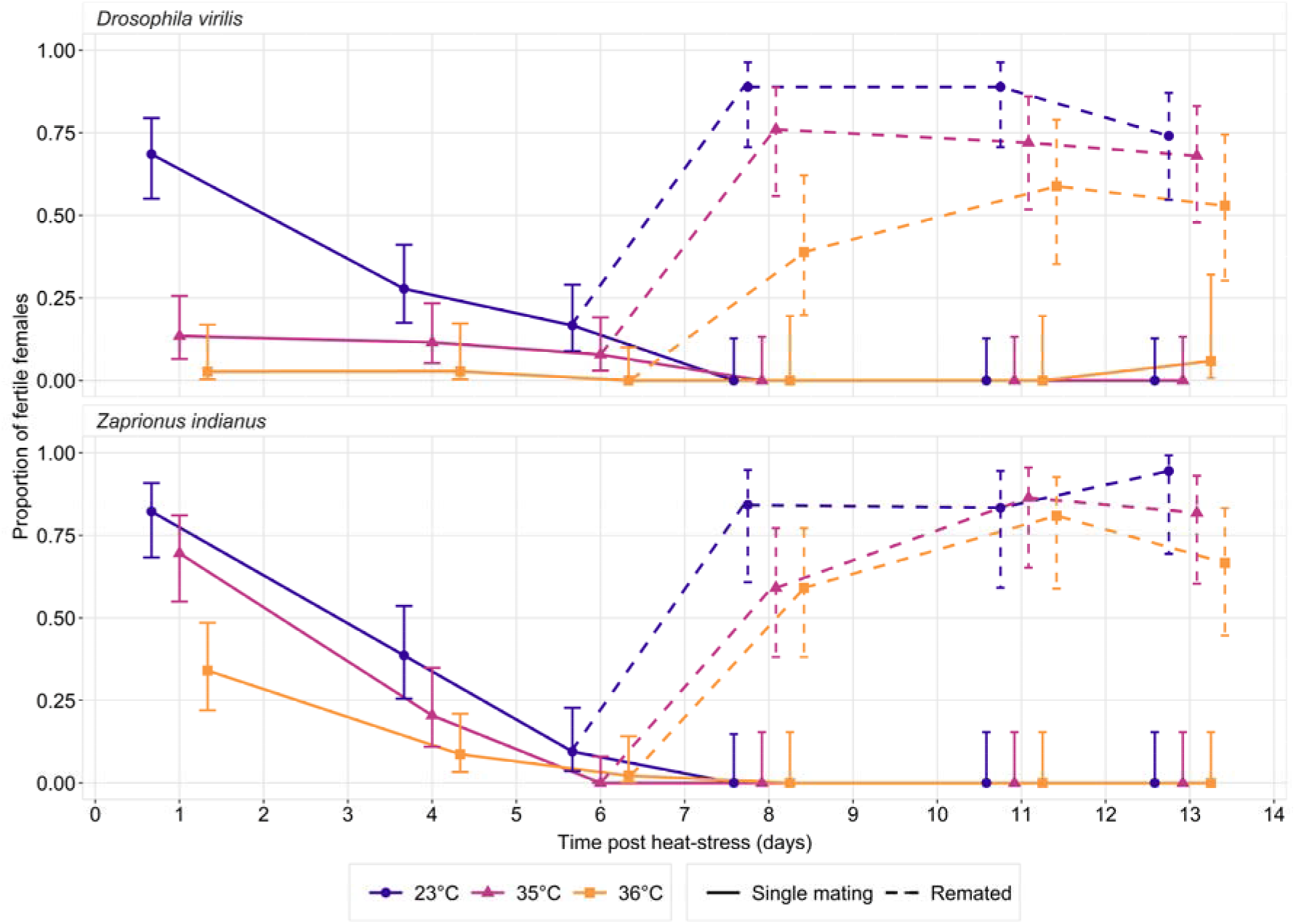
Proportion of fertile *D. virilis and Z. indianus* females over time for **Experiment 2: Mated/Heat/Isolated** as described in Fig 1. Mated females were heat-shocked at either benign (23°) or two stress temperatures (35 & 36°C) for 4 hours. Following heat stress, all females were isolated and allowed to lay eggs in fresh vials three times. After 6 days, the experiment was split into two treatments. **2a Mated/Heat/FullyIsolated:** females remained isolated and moved onto three fresh vials to lay any remaining eggs. **2b Mated/Heat/Isolated/Remated:** focal females were paired with new male partners, and the mating group were given 3 fresh vials to produce offspring. Error bars represent 95% confidence intervals.

There was a significant interaction between temperature and time on fertility of *D. virilis* in Experiment 2 prior to treatment splitting (χ^2^_(2)_ = 9.943, p<0.007; Figure 3). Fertility of controls started high immediately following heat treatment and fell over time, whereas fertility at stress temperatures started low and remained low for the duration. There was a main effect of temperature (χ^2^_(2)_ =21.146, p<0.001; Figure 3) and time (χ^2^_(1)_ =17.352, p<0.001; Figure 3) on fertility of *D. virilis* in Experiment 2. Both stress temperatures showed lower fertility than controls, and all treatments showed a decline in fertility over time.

There was no significant interaction between temperature and time on fertility of *Z. indianus* from Experiment 2 (χ^2^_(2)_ = 1.777, p=0.411; Figure 3). However, there was a significant overall effect of temperature (χ^2^_(2)_ =80.161, p<0.001; Figure 3) and time (χ^2^_(1)_ =99.756, p<0.001; Figure 3) on fertility of *Z. indianus*. In this species the highest temperature of 36°C results in significantly lower fertility than both controls (p<0.001) and the stress temperature of 35°C (p<0.001). All temperatures result in a loss of fertility over time.

### Experiment 2b: Mated/Heat/Isolated/Remated

There was no significant interaction between temperature and time on fertility of *D. virilise* after females were given the chance to remate in Experiment 2b (χ^2^_(2)_ = 3.549, p=0.170; Figure 3). However, we found a significant effect of temperature on fertility in *D. virilis* (χ^2^_(2)_ =9.520, p=0.009; Figure 3). Specifically, fertility of females exposed to the stress temperature of 36°C was significantly lower than fertility from the control 23°C (p=0.002) and stress temperature of 35°C (p=0.046). There was no significant effect of time on fertility (χ^2^_(1)_ =0.515, p=0.473; Figure 3).

There was no significant interaction between temperature and time on fertility of *Z. indianus* when females were given the opportunity to remate in Experiment 2b (χ^2^_(2)_ = 1.049, p=0.592; Figure 3). There was also no main effect of temperature on fertility (χ^2^_(2)_ =4.250, p=0.119; Figure 3). However, there was a significant effect of time on fertility (χ^2^_(1)_ =4.775, p=0.029; Figure 3), where fertility slightly increases over time.

## Discussion

We found little evidence that virgin females are susceptible to fertility loss at high temperatures. Heat-stress did not influence fertility of virgin *D. virilis* or *Z. indianus* females that were then mated after heat-stress. Fertility of *Z. indianus* females was initially lower at the first time-point measured post heat-stress, and increased over the duration of the experiment. Conversely, fertility of *D. virilis* females was consistently high over the duration, suggesting that *Z. indianus* females were slower to mate and produce offspring with their paired males than *D. virilis*.

Mated females given no opportunity to remate used up their viable sperm reserves within the first week of laying. However, we found that heat stress sterilised females of both species, likely through destruction of stored mature sperm. This is curious because mature sperm in males of both species appear to be largely unaffected by the same temperature treatments (Parratt et al., 2021). We find that mated females are sterilised at temperatures around 2°C lower than those required to completely sterilise 80% of males from our study species (Parratt et al., 2021). Hence our results suggest that females of both species are worse at protecting mature sperm from high temperatures than males.

We found that the temperatures required to sterilise mated females differ between the two species. Four hours at either 35°C or 36°C almost completely sterilise *D. virilis* females (∼90% of females produce no offspring), whereas mated *Z. indianus* females are mostly fertile when stressed at 35°C and only a small majority are sterilised when exposed to 36°C for four hours (∼60% of females produce no offspring). The finding that mature sperm from *Z. indianus* is likely more resilient than sperm from *D. virilis* is consistent with our previous study that heated adult males of each species. Males of *D. virilis* require temperatures of no less than 37°C for 4h to immediately sterilise the majority of males, whereas males of *Z. indianus* are fertile up to their lethal temperature of ∼38°C (Parratt et al., 2021). While the absolute temperatures required to sterilise males and mated females are different, these results combine to suggest that mature sperm from *Z. indianus* are generally more thermally robust than those from *D. virilis*. It is unclear exactly why this may be the case, however *Z. indianus* tend to live in slightly warmer areas than *D. virilis*. The temperature experienced by individuals at the upper edge of their thermal range in the hottest month of the year (Tmax+1sd: WorldClim.org BIO05) is 36.1°C for *Z. indianus*, whereas it is 32.6°C in *D. virilis* (Parratt et al., 2021). Therefore, *Z. indianus* sperm may better adapted to high temperatures than *D. virilis*, although this is beyond the scope of this study.

To unpick effects of high temperatures on stored sperm from direct effects on females, we then gave a chance for mated females to ‘recover’ fertility after they had used up their viable stored sperm. We found that while the majority of females exposed to all temperatures were able to produce offspring when paired with new males, females heated at 36°C performed worse than controls in *D. virilis*. Therefore, it is likely that 36°C thermal stress results in some permanent damage to females of this species. However, the almost complete sterilisation of sperm stored in female *D. virilis* paired with a general capacity to ‘recover’ fertility suggests that initial sterilisation in this species is likely due to the destruction of stored sperm by high temperatures and not direct effects on females. Mated *Z. indianus* females were equally able to recover fertility when paired with new males, regardless of the heat-stress temperature experienced. While the temperatures required to reduce fertility of mated females were higher in this species, there was no long-term effects of temperature on female recovery when females were presented with new males, suggesting that this initial reduction of fertility in *Z. indianus* is also driven by effects on stored sperm.

Sterilisation of mated females could be particularly devastating to species with low remating rates. However, females can use facultative polyandry to improve offspring production when mating with sub-fertile males (Sutter et al., 2019; Vasudeva et al., 2021). For example, heat-shocked males of the flour beetle *Tribolium castaneum* have low numbers of viable sperm after heat-stress (Vasudeva et al., 2021). Here, females increase their remating rate when mated with a heat-shocked male, rescuing fertility to normal levels. However, whether increased polyandry is observed when sperm within the female is sterilised by high temperatures remains an open question. Also, there may be species where facultative polyandry is impossible, for example in seasonally reproducing animals with discrete mating opportunities. Those particularly at risk include species that store sperm for long periods of time, such as hymenopteran insects that have been observed to store sperm for up to 10 years (Keller, 1998; Pamilo, 1991). In these cases, sterilisation of mated females may actually be worse for population persistence than sterilisation of males.

Understanding how high temperatures affect male fertility has improved our ability to predict the consequences of climate change on species (Parratt et al., 2021; van Heerwaarden and Sgrò, 2021; Walsh et al., 2019b). When these severe long-term effects on male fertility are combined with the immediate sterilisation of mated females like we have demonstrated, the impact of rising temperatures on wild populations may be exacerbated. Further, we find here that the temperatures required to sterilise mated females are not always consistent with the temperatures required to sterilise males. It will be important to determine whether this is true across species and taxa to help forecast vulnerability climate warming effects. Species where sperm in both males and mated females cannot be protected may be particularly vulnerable, whereas species where females can protect sperm effectively may be more resilient to an increasing incidence and severity of heat-waves.

## Acknowledgements

The authors thank Jolanta Tanianis-Hughes for assistance with experiments and Sophie Lyth for helpful advice regarding experimental design.

## Data and materials availability

All data and analysis R code is available at https://datadryad.org/stash/share/7wn67Q4UVZBXStL1OKTk87xJ9CzXh-GrQ1H2ZoxC7TA

